# Immunoengineered Nanobead System for the Isolation and Detection of Circulating Tumor Cells

**DOI:** 10.1101/2021.01.18.427201

**Authors:** Pengfei Zhang, Mohamed S. Draz, Anwen Xiong, Wannian Yan, Huanxing Han, Wansheng Chen

**Affiliations:** Department of Pharmacy, Department of Laboratory Medicine, Changzheng Hospital, the Second Military Medical University, Shanghai 200433, China; Department of Central Laboratory, Shanghai Skin Disease Hospital, Tongji University School of Medicine, Shanghai 200443, China; Department of Chemistry and Chemical Biology, Harvard University, Cambridge, Massachusetts, USA; Wyss Institute for Biologically Inspired Engineering, Harvard University, Cambridge, Massachusetts, USA; Department of Medical Oncology, Shanghai Pulmonary Hospital, Tongji University Medical School Cancer Institute, Tongji University School of Medicine, Shanghai 200433, China; Ailex Technology Group Co., Ltd. Shanghai 201108, China

**Keywords:** magnetic nanoparticle, quantum dots, fluorescent magnetic nanobeads, circulating tumor cells, simultaneous capture and detection

## Abstract

Highly efficient capture and detection of circulating tumor cells (CTCs) remain elusive mainly because of their extremely low concentration in the peripheral blood of patients. Herein, we present an approach for the simultaneous capturing, isolation, and detection of CTCs using an immuno-fluorescent magnetic nanobead system (iFMNS) coated with a monoclonal anti-EpCAM antibody. The developed antibody nanobead system allows magnetic isolation and fluorescent-based quantification of CTCs. The expression of EpCAM on the surface of captured CTCs could be directly visualized without additional immune-fluorescent labeling. Our approach is shown to result in a 70 - 95% capture efficiency of CTCs, and 95% of the captured cells remain viable. Using our approach, the isolated cells could be directly used for culture, reverse transcription-polymerase chain reaction (RT-PCR), and immunocytochemistry (ICC) identification. We applied iFMNS for testing CTCs in peripheral blood samples from a lung cancer patient, which suggested that this approach would be a promising tool for CTCs enrichment and detection in one step.

## Introduction

Circulating tumor cells (CTCs) are free tumor cells shed from original or metastatic tumors into the peripheral blood. CTCs play critical roles in cancer metastasis, resulting in 90% of cancer-related deaths[1–3]. Capturing, characterization, and enumeration of CTCs were considered prognostic biomarkers in tumor metastasis and cancer diagnostics [4, 5]. However, the limited sensitivity of commercially available approaches combined with the disease’s complexity and heterogeneity had restricted the broad acceptance and dissemination of CTC-based diagnostics. Thus, highly efficient isolation and analysis of CTCs from the whole blood is an urgent clinical need.

Immunomagnetic separation represents a promising approach for cell isolation because of its capability to rapidly process a large volume of samples and easy operation and facile cell recovery by the end users[6, 7]. The Cell Search System[8, 9], a U. S. Food and Drug Administration approved platform for clinical CTC enrichment, provides low capture purities (<1%) with high background of white blood cells. Recently, some analytical methods with different signaling modes have been explored to detect CTCs, including electrochemistry[10–12], inductively coupled plasma-mass spectrometry[13], Raman imaging[14, 15], colorimetry[16, 17], and fluorescence[18]. Among these detection techniques, fluorescence-based assays have attracted much interest due to the quick response, high sensitivity, non-destructivity, and real-time monitoring. Many organic fluorescent dyes and fluorescent nanoprobes, such as upconversion nanoparticles[19, 20], quantum dots [21, 22] and gold nanoclusters with visible light emission[23–25], have been applied for the detection of biological targets circulating in blood including CTCs[26–28].

An ideal probe for CTCs measurement would allow magnetic isolation and fluorescent-based detection of the cells. Such probes require a compact structure with a rapid magnetic response, high specificity and minimal nonspecific binding, and strong fluorescent signals for CTCs detection and quantification. Quantum dots (QDs) with unique optical properties, including tunable wavelength, high quantum yields, and photobleaching resistance, are ideal for the preparation of fluorescent magnetic nanoprobes. QDs based fluorescent magnetic nanobeads (FMN) prepared by swelling with polymer nanospheres[29, 30], layer-by-layer self-assembly[31–33], DNA templated hybridization[21, 34, 35], silica shell coating, or polymer assembly[36], have been reported for CTCs isolation and identification. However, these probes still suffer from complicated preparation steps, low fluorescent signals, and weak magnetic response.

Here, a fluorescent magnetic nanobeads system (iFMNS) with a highly bright fluorescent intensity that allows immunomagnetic separation of CTCs was prepared via a simple emulsion/evaporation method. The prepared FMN has a diameter of 114 nm with considerable colloidal stability (i.e., did not aggregate or precipitate in the buffer during incubation or centrifugation,) and ultrabright fluorescence (encapsulated with many QDs) can improve the fluorescent immunoassay sensitivity compared with single QDs immunoprobe[37]. Together with these unique properties, the anti-EpCAM antibody was conjugated with FMN via streptavidin-biotin bridges. The obtained anti-EpCAM antibody-modified FMN nanoprobes were successfully used for cancer cell isolation with efficiency in a range of 70-95% of the tested cells within 10 min using an external magnetic field. The cells isolated by this method are directly identified by the fluorescent signals generated from iFMNS that exist on the surface of isolated CTCs without any additional immunofluorescent labeling procedures. These small biocompatible nanobeads have nearly no influence on the viability and propagation of CTCs, which is of significant importance for testing the isolated cells using cellular and molecular analysis techniques. Furthermore, this method was successfully validated on a lung cancer patient peripheral blood sample and showed potential for clinical application.

## Materials and methods

### Materials

Oil-soluble CdSe/ZnS QDs were obtained from Suzhou Xingshuo Nanotech. Oleic acid capped magnetic nanoparticles (MNPs) with a diameter of 10 ± 5 nm were purchased from Nanjing Nanoeast Biotech. Poly(styrene-co-maleic anhydride) (PSMA) copolymer, streptavidin, bovine serum albumin (BSA), N-(3-(dimethylamino)propyl)-N’-ethylcarbodiimide hydrochloride (EDC), were obtained from Sigma-Aldrich. The biotinylated anti-EpCAM antibody was obtained from R&D Systems. Red blood cell (RBC) lysis buffer was bought from Sangon Biotech. FITC anti-mouse/human CD45 antibody(FITC-CD45) was purchased from BioLegend. Blood samples were obtained from Shanghai Changzheng Hospital.

Human cell lines of gastric cancer cells (SGC-7901), pancreas cancer cells (PANC1), lymphoma cells (Raji), EGFR-mutated (HCC827), and non-EGFR-mutated (A549) lung cancer cells were obtained from the Cell Bank of the Chinese Academy of Sciences. All the cell lines were cultured in Dulbecco’s modified Eagle’s medium (DMEM) medium except Raji cells in RPMI-1640 medium, containing 10% fetal bovine serum at 37 °C in a humidified 5% CO_2_ atmosphere.

### Fabrication of FMN

Fluorescent magnetic nanobeads (FMN) were prepared according to our previously reported method with a slight modification[37]. Typically, one milliliter of a chloroform solution containing PSMA (10 mg), QDs (10 mg), and MNPs (10 mg) was added into a four mL aqueous solution containing sodium dodecyl sulfate (11.5 mg). After magnetic stirring for 3 min at 300 rpm, the miniemulsion was prepared through ultrasonication for 100 s (10 s pulse, 10 s pause) at 50% amplitude with a sonicator (Fisher Sonic Dismembrator Model 500, USA) under ice-bath cooling to prevent the evaporation of chloroform. Afterward, the prepared miniemulsion was magnetically stirred at 300 rpm overnight in an open container to evaporate chloroform. The resulting nanobeads with PSMA as matrix polymer were pelleted at 12,000 rpm for 10 min, and the supernatant was discarded. The pellet of nanobeads was washed with deionized water three times. After that, the FMN was purified by magnetic separation and stored in deionized water at 4 °C in the dark for further use.

### Conjugation of the anti-EpCAM antibody with FMN

Firstly, the covalent conjugation of streptavidin (SA) onto FMN was achieved by the classic EDC coupling reaction of carboxyl groups on the surface of FMN and amino groups streptavidin. Typically, FMN (1 mg) in 400 μL of phosphate buffer (25mM, pH 6.0), and EDC (500 nmol) were added into a tube, and incubated for 30 min to activate carboxyl groups. Excess crosslinkers were removed by centrifugation at 12,000 rpm for 10 min and decantation of supernatant. Then, SA (60μg) was added into activated FMN, and the mixture was incubated at room temperature for 2 h. To block unreacted carboxyl groups on the surface of FMN, BSA (0.5 mg) was added into the tube and incubated for another 30 min. The mixture was purified by centrifugation at 10,000 rpm for 0.5 h and washed with PBST buffer (PBS with 0.05% Tween-20) twice to remove free SA molecules. The purified FMN-SA conjugates were dispersed in PBS buffer with 1% BSA and stored at 4 °C for further use.

Anti-EpCAM antibody conjugated iFMNS were freshly prepared just before the cell separation. Typically, 0.1 mg FMN-SA and 10 μg biotinylated anti-EpCAM incubated in PBS buffer for 0.5 h at room temperature. Then anti-EpCAM conjugated iFMNS were obtained by magnetic separation and used for cell separation.

### Capturing of tumor cells

A defined number of cancer cells were suspended in PBS buffer or whole human blood. To capture the cancer cells, iFMNS were first incubated with a cell suspension for 10 min, followed by removing unbound iFMNS through centrifugation at 800 rpm for 3 min. The resulting cell suspension was collected by applying magnetic columns separator (Miltenyi, Germany) and washed twice with PBS buffer. The numbers of captured cells were counted using a hemocytometer through direct counting with bright light microscopy. The tumor cells were pre-labeled using green cell tracker dye (CellTrace CFSE, Thermo Fisher, USA), which could be distinguished from the red FMN, before the capturing testing experiment. The captured cells were observed using a fluorescent microscope (BX-51 Olympus, Japan). Capture efficiency was calculated using the following Equation.

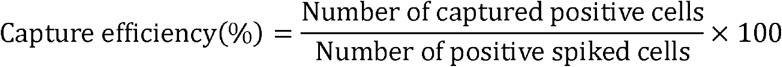

### Identification of tumor cells

After magnetic separation, the captured cells were plated in a closed chamber drawing by a super pap pen on a clean glass slide, fixed with 4% paraformaldehyde for 10 min, permeabilized with 0.1%Triton-X 100 for 10 min, blocked with 3% BSA for 30 min, and stained with FITC-CD45 antibody, and 20 μg/mL DAPI for 10min. Cells that were EpCAM^+^/DAPI^+^/CD45^−^ were scored as CTCs.

### Cell viability analysis

The cell dead/live of the isolated cells was analyzed by staining dead cells with trypan blue, while live cells exclude dying. Analysis of cell apoptosis was performed using Annexin V-FITC/PI staining assay, and the cells were subsequently counted by flow cytometry (CyAn ADP, Beckman Coulter, USA). Further, SGC-7901 cells captured with FMN were cultured at 37 °C with DMEM containing 10% fetal bovine serum, 60 μg/mL penicillin G, and 100 μg/mL streptomycin sulfate in a humidified atmosphere with 5% CO_2_. Sterile conditions should be maintained during the whole procedure.

### EGFR mutation analysis of captured cells

Spiked in HCC827 and A549 lung cancer cells in blood samples were magnetically enriched by the iFMNS. Total RNA was extracted from cells using the TRIzol reagent (Thermo Fisher, USA). For mRNA studies, cDNA using PrimeScript™ RT Master Mix (Takara) and quantitative PCR (qPCR) was performed to detect EGFR levels by using SYBR-Green dye (Takara). Further, the qPCR analysis was performed on a quantitative PCR machine (ABI-7900HT, USA) using an amplification process of initial denaturation at 95 °C for 2 min, 40 cycles at 95 °C for 30s, 58 °C for 30s, and 72 °C for 30s. Relative expressions were calculated using the 2^-∆∆Ct^ method and GAPDH was served as a loading control. Primers for EGFR mutational analysis were designed from the EGFR 21 exon gene sequence, using Primer premier 5.0 software. And the primers of GAPDH were also created using the premier 5.0 software. Primers were obtained from Shanghai Sangon Biotech. A pair of primer (Forward primer: 5’-TTCCCATGATGATCTGTCCCTC-3’; Reverse primer: 5’-CACCTCCTTACTTTGCCT-3’) could amplify a 214 bp product of EGFR exon 21 sequences. The primers for GAPDH were as follows: Forward primer: 5’-AATCCCATCACCATCTTC-3’; Reverse primer: 5’-AGGCTGTTGTCATACTTC-3’. The qPCR products were separated in a 2% (wt/vol) agarose gel.

### Detection of CTCs in Cancer Patient Peripheral Blood Samples

Blood samples from a lung cancer patient and healthy normal controls were collected and detected with iFMNS. In each typical assay, a certain amount of blood was incubated with iFMNS for 10 min. After enrichment, the isolated cells were incubated with the FITC-CD45 and DAPI and observed by a fluorescence microscope, as previously described in Section: Identification of Tumor Cells. The cells with phenotypes of EpCAM positive and DAPI positive but CD45 negative were enumerated as CTCs. The Ethics Committee approved Studystudy and data analyses of Shanghai Changzheng Hospital. Patient consent was not obtained as all personal identifiers, and patient information was delinked from the serum specimens.

## Results and discussion

### Preparation and characterization of FMN

Fluorescent magnetic nanobeads were prepared by an oil-in-water emulsion-evaporation technique (**Figure 1**). In this approach, poly(styrene-co-maleic anhydride) (PSMA) was selected as the matrix material for nanobeads because PSMA is initially oil-soluble and then can be transferred into a hydrophilic surface by hydrolysis of anhydride groups into carboxyl groups after nanobeads formation. Oil-soluble QDs and MNPs dispersed in PSMA polymer chloroform solution used as the oil phase. Subsequently, an oil/water miniemulsion was formed through stirring and ultrasonication. The gradual removal of chloroform from the aqueous phase by evaporation was compensated by transferring chloroform from the mini droplets to the aqueous phase. Finally, the polymer precipitated, entrapped QDs and MNPs inside polymer nanobeads, and formed polymer/nanoparticle hybrid nanobeads.

**Figure 1.**
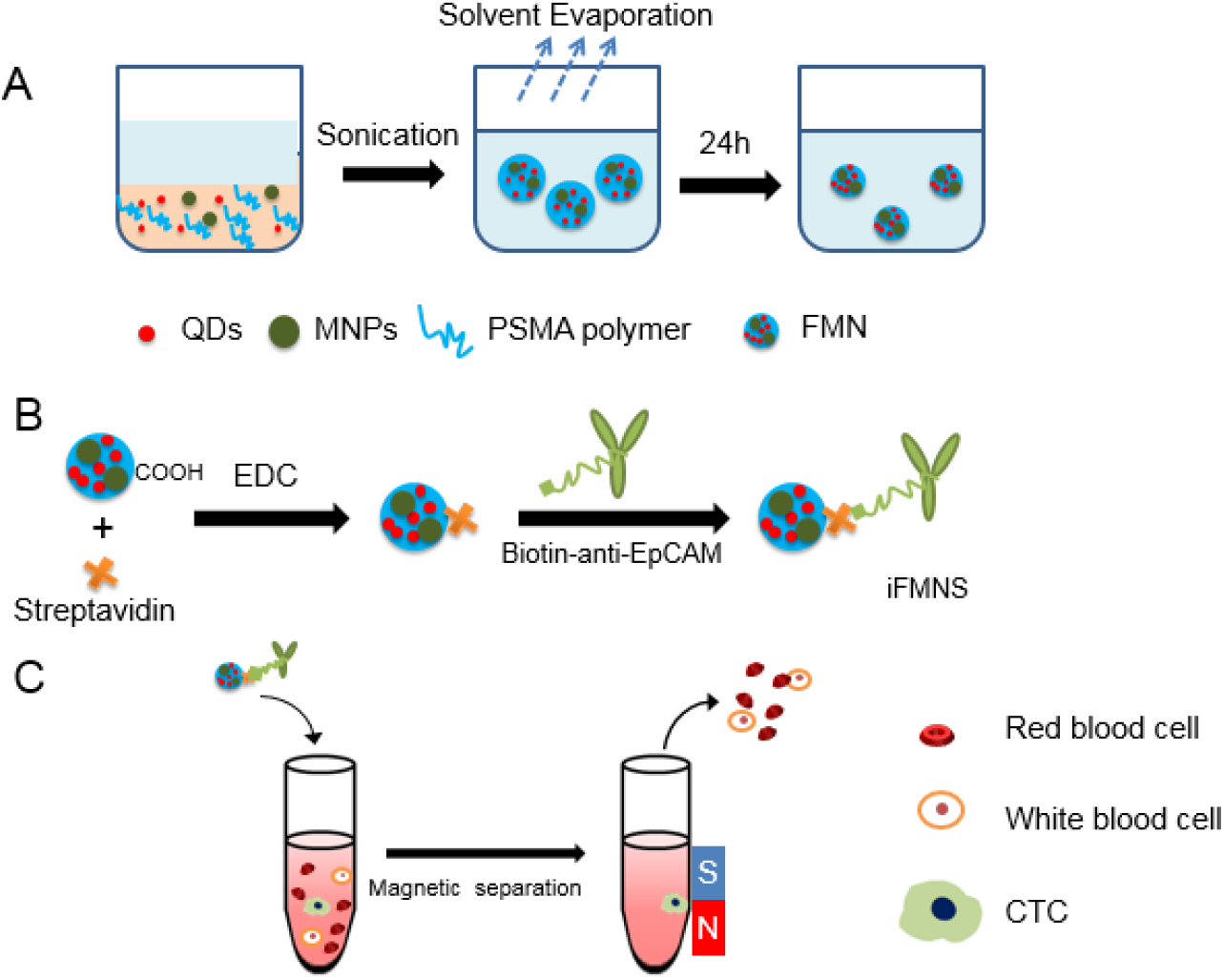
Schematic illumination of CTC capturing using iFMNS. (A) Encapsulation of QDs and MNPs into PSMA nanobeads by miniemulsion and solvent evaporation techniques. (B) Construction of iFMNS conjugate. Carboxyl capped FMN and streptavidin were covalently conjugated using EDC crosslinker, and biotinylated anti-EpCAM antibody was immobilized to the streptavidin-coated FMN. (C) Specific recognition and isolation of CTCs by iFMNS.

The TEM image of the prepared FMN, as shown in Figure 2A, demonstrates that the FMN has a diameter in the range of about 40-95 nm with a broad size distribution. However, using the dynamic light scattering technique, the average hydrodynamic diameter of FMN was 114 nm and a polydispersity index (PDI) of 0.13 (Figure 2B), indicating the relatively monodisperse and stability of the prepared system. Due to the mild reaction process and the polymer matrix isolation between QDs and MNPs, QDs encapsulated in the polymer nanospheres still possess high fluorescent intensity. The digital image of FMN suspension (Figure 2C) was highly fluorescent under UV light illumination. It can be rapidly isolated (in 10 min) from the solution to the tube wall using an external magnet.

**Figure 2.**
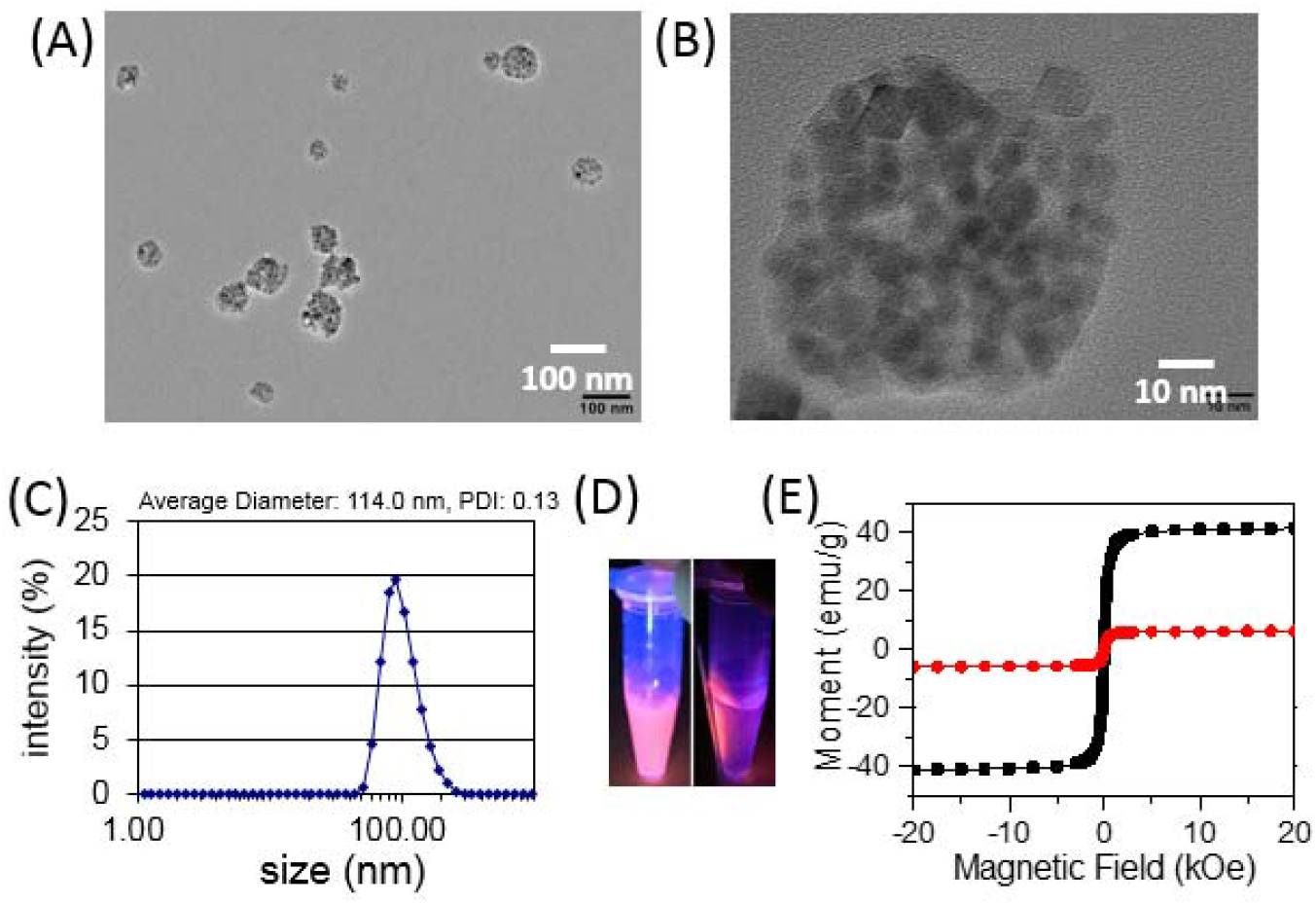
(A) TEM images of FMN. (B) Hydrodynamic diameters and PDI of the FMN. (C) Photographs of FMN solution pre and post-isolation using a magnet under the UV lamp excitation. (D)Magnetic hysteresis loop of the FMN encapsulated with different concentrations of MNPs measured at room temperature.

From the magnetic hysteresis loop characterization (Figure 2D), the FMN had an excellent superparamagnetic property at room temperature with a tunable magnetic saturation value (41.6 emu/g and 6.04 emu/g) by adjusting the concentration of magnetic nanoparticles added in the oil phase. The high magnetic saturation FMN was used for the construction of iFMNS. To apply the FMN for CTCs isolation and detection, as shown in Figure1, streptavidin was covalently coated on the surface of FMN by a classic carboxyl-amino reaction under the activation of carbodiimide. Then biotinylated anti-EpCAM antibody was attached to the surface by the streptavidin-biotin interaction just before cell isolation.

### Tumor cell capturing using iFMNS

The iFMNS (FMN + anti-EpCAM antibody) concentration applied for cell capture was preoptimized using a suspension of SGC-7901 cells. The results are shown in Figure 3A. With the increase of iFMNS concentration, the capture efficiency increased until the concentration reached 0.2 mg/mL, and 96% of SGC-7901cells were captured. Then, the magnetic attraction time was investigated, as shown in Figure 3B, nearly 100% of tumor cells were captured in approximately 8 min. Finally, we used 0.3 mg/mL iFMNS under 10 min of magnetic isolation for further experiments. The magnetic isolation of cells was respectively performed using FMN-SA and iFMNS as capturing probes. As shown in Figure 3C, nearly no cancer cells (3%) were captured by FMN-SA indicated that our iFMNS (with anti-EpCAM antibody) was specifically binding to cancer cells. Moreover, the iFMNS were also applied for several other cancer cell line isolation, as shown in Figure 3D. The EpCAM-negative lymphoblast Raji cells were not isolated by magnetic separation, which also indicated the specificity of the prepared iFMNS. The cell capture efficiencies were more than 70% in various kinds of cancer cells, showing that iFMNS had general applicability for EpCAM-positive cancer cells.

**Figure 3.**
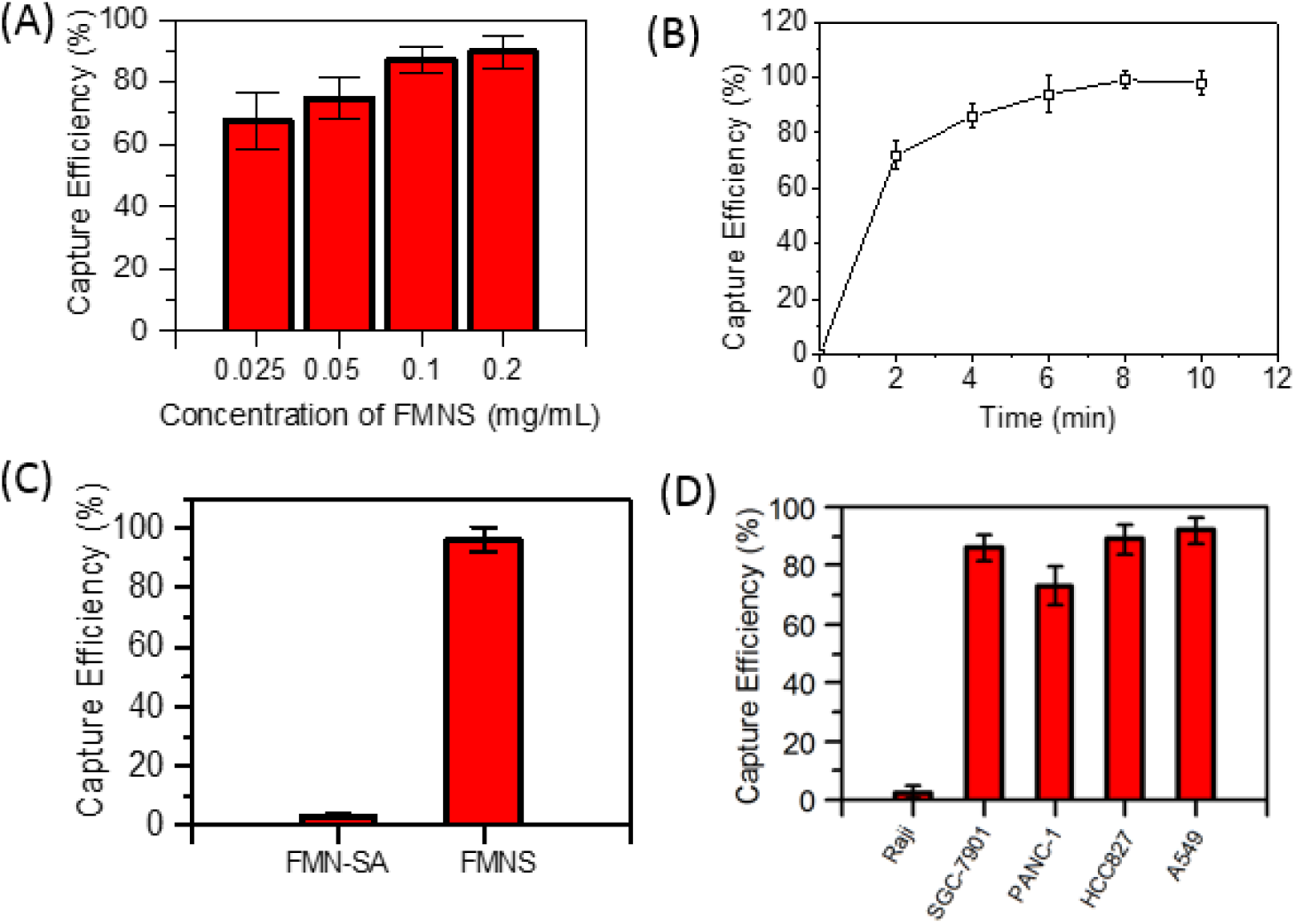
(A) Capture Efficiencies ofSGC-7901 cells in PBS buffer with different concentrations of iFMNS. (B) Capture efficiencies of SGC-7901 cells using iFMNS by a magnet with different attraction time. (C) Capture efficiencies of SGC-7901 cells in PBS buffer with FMN-SA and iFMNS nanoprobes. (D) Capture efficiencies of iFMNS to Raji, SGC-7901, PANC-1 cells, HCC827, and A549 tumor cell lines.

Then, the capability of iFMNS to capture tumor cells to mimic CTC blood samples was investigated. The samples were prepared by spiking green cell trace dye pre-stained SGC-7901 cells into whole blood. The captured cells were enumerated under a fluorescence microscope. The tumor cells and red blood cells were effectively separated from each other using magnet separation in a centrifuge tube, as shown in Figure 4A. Although there were still some red blood cells in the isolated tumor cell suspension mainly due to the gravitational sedimentation, the isolation impurity can be improved primarily by using a magnetic column.

**Figure 4.**
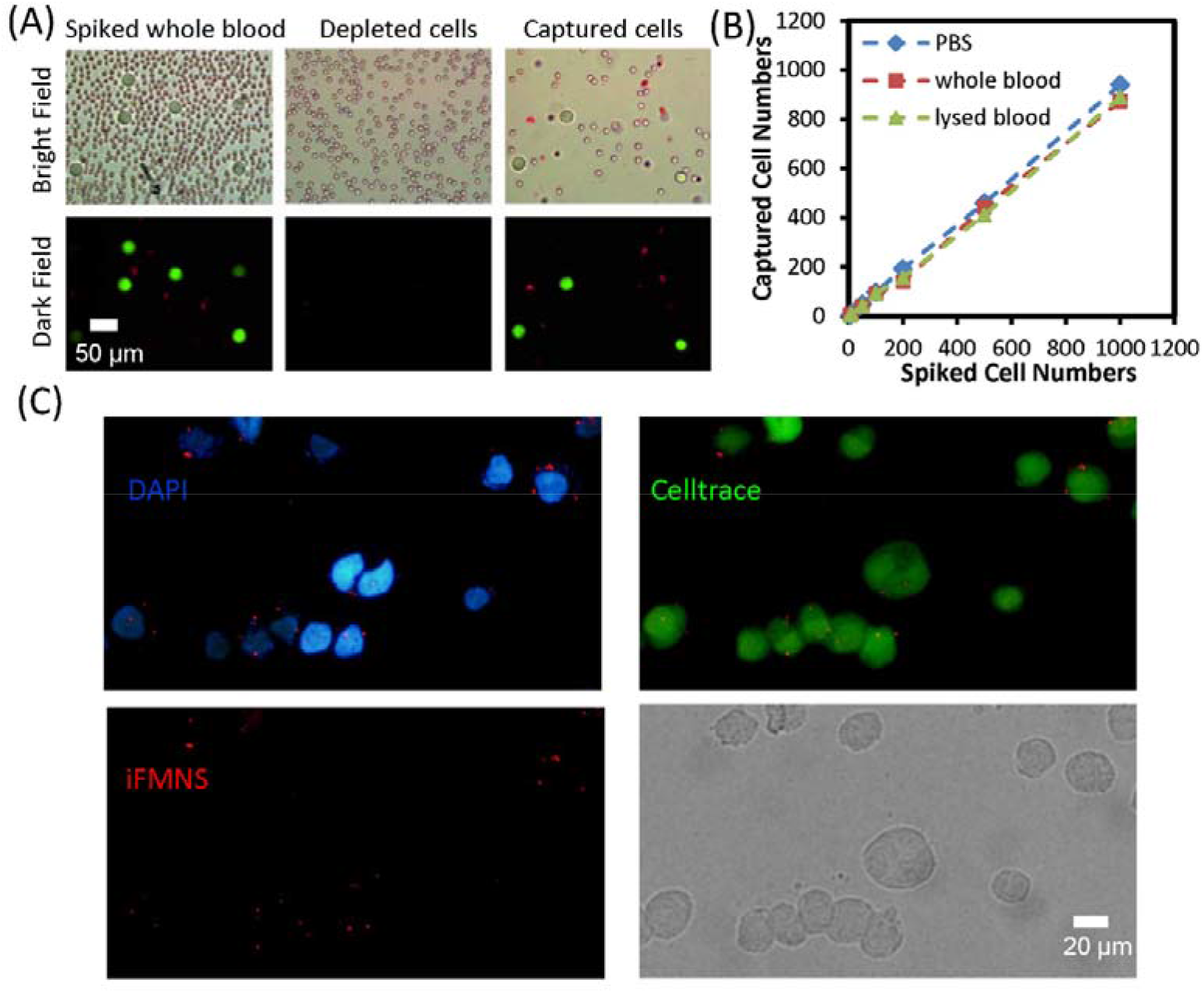
(A) Microscopic images of the initial mimic clinical sample, the depleted cells from the mimic clinical sample, and the enriched cells (arrow: SGC-7901 cells). (B) Capture efficiencies with iFMNS at different cell concentrations (5-1000 cells/mL) in three different types of samples: PBS, whole blood, and lysed blood. (C) Fluorescence microscope images of Celltrace pre-labeled SGC-7901 tumor cells captured by iFMNS (red) and DAPI (blue) stained-cells.

For comparison of the matrix effect, CTC capture efficiencies were also examined in PBS, whole blood, and lysed blood spiked with CTC at concentrations of5-1000 cells/mL. The results are shown in Figure 4B. The capture efficiencies in the three types of samples were comparable and did not have significant differences. These results suggested that complex conditions had negligible effects on the binding between iFMNS and the target cells, and iFMNS could be directly used in whole blood.

### Identification of the captured cells

The isolated CTCs were identified based on fluorescent labeling and morphology. With the ultra-bright fluorescent intensity of iFMNS and quick response magnetic separation performance, we investigate the specific recognition and labeling of iFMNS towards tumor cells. As shown in Figure 4C, the green cell trace pre-stained SGC-7901 cells were specifically isolated. The EpCAM motif on the surface of SGC-7901 cells can be directly visualized by the red fluorescent signals from the as-prepared iFMNS. While the different expression amount of EpCAM can also be semi-quantified through fluorescent intensity from iFMNS on the cell surface.

Besides, FITC labeled anti-CD45 (CD45-FITC, green) stained with WBCs and DAPI (blue) stained nuclear. The three-color was used to discriminate tumor cells from WBCs. CTCs are defined as a positive signal for EpCAM (iFMNS, red), DAPI stained nuclear and negative signal for CD45-FITC, while WBCs are defined as a negative signal for iFMNS, positive signal for DAPI, and CD45-FITC. Figure 5 shows that the intense red and blue fluorescence were observed within DAPI and iFMNS stained SGC-7901 cells. Meanwhile, no fluorescence signal was observed for the channel of CD45-FITC. As a control, WBC cells, which are CD45 positive and EpCAM negative, are easily stained with green fluorescence but without red fluorescence. These observations describe that CTCs are specifically recognized in the blood samples from WBCs. The results confirm that the as-constructed system could efficiently isolate and identify CTCs to mimic clinical samples.

**Figure 5.**
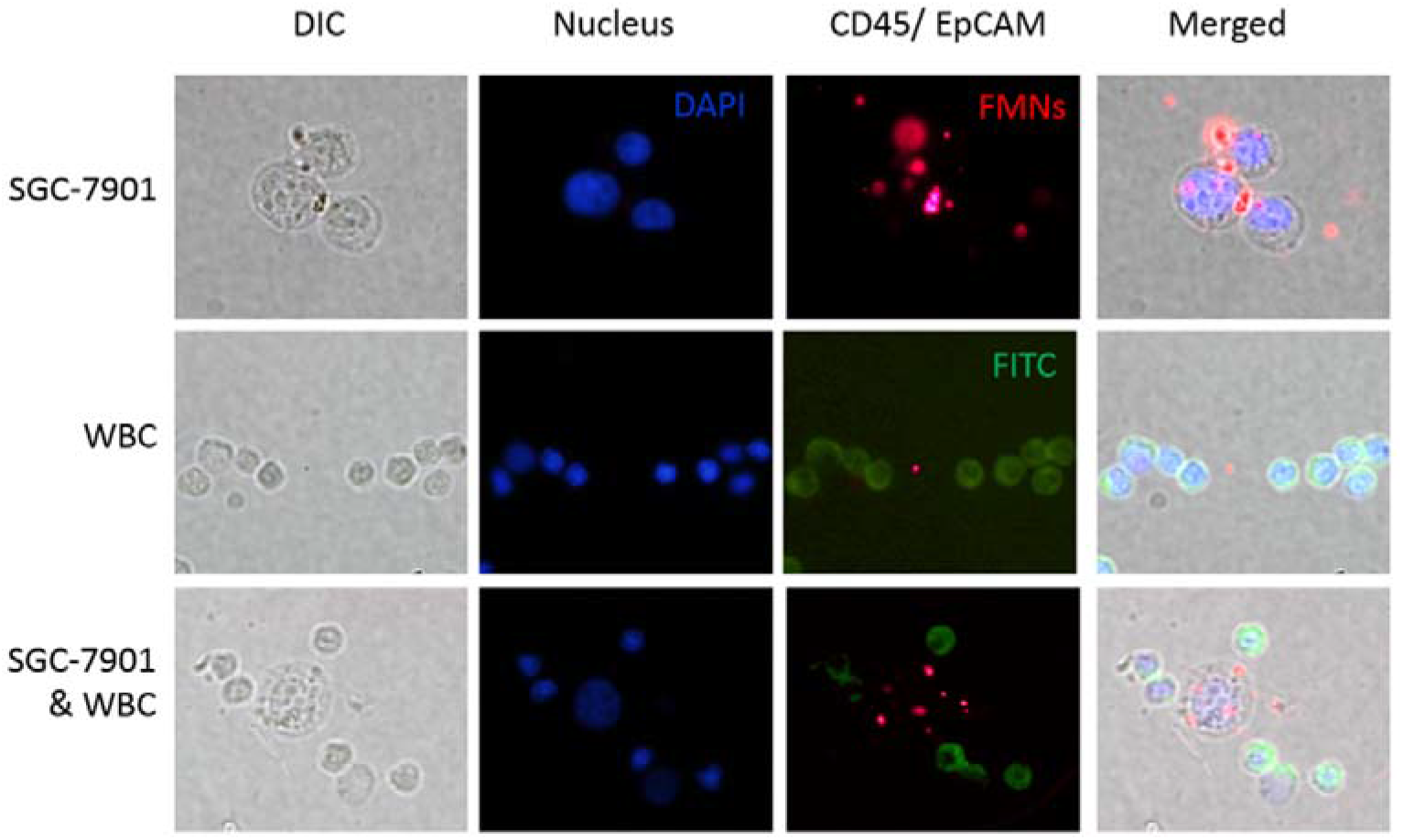
Microscopic images of SGC-7901 cells captured from mimic clinical blood samples and identified with the three-color fluorescent labeling. Nucleus (DAPI): excitation 405 nm, emission 447 nm. CD45 (FITC): excitation 488 nm, emission 525 nm. EpCAM (iFMNS): excitation 488nm, emission 620 nm. Merged: merge of the nucleus (DAPI), CD45 (FITC), and EpCAM (iFMNS).

### Cell viability and function of the captured cells

The viability of isolated tumor cells was investigated to evaluate this isolation method’s influence on the cell for further culture and analysis. The viability of cancer cells was measured by trypan blue dyeing. As shown in Figure 6A, the dead cells would be stained by trypan blue dyes, while the live cells would not be stained. The viability rate of isolated tumor cells was calculated to be about 95%, indicating that most of the tumor cells are live after magnetic isolation. Apoptosis analysis of tumor cells before and after magnetic isolation was performed by FITC-annexin V/PI staining and flow cytometry. As shown in Figure 6B, the proportion of apoptotic cells, including early and late, for the control group was 9.75%, while for captured tumor cells was 5.97%. These results indicated that magnetic separation using iFMNS would not induce the tumor cells apoptosis. The slightly lower proportion of apoptosis in captured tumor cells may attribute to the low expression of EpCAM in apoptosis tumor cells, which would not be captured by iFMNS. Then, the captured SGC-7901 tumor cells were re-cultured and propagated *in vitro*. As shown in Figure 6C, the iFMNS captured tumor cells proliferated without a significant change in behavior and morphology compared with the control samples of SGC-7901 cell lines. Interestingly, cadmium-containing quantum dots are limited in biological labeling mainly due to toxicity for cell growth, but as-prepared iFMNS had nearly no influence on the cell growth and apoptosis be attributed to the thick polymer shell encapsulation around QDs to prevent cadmium ions leakage.

**Figure 6.**
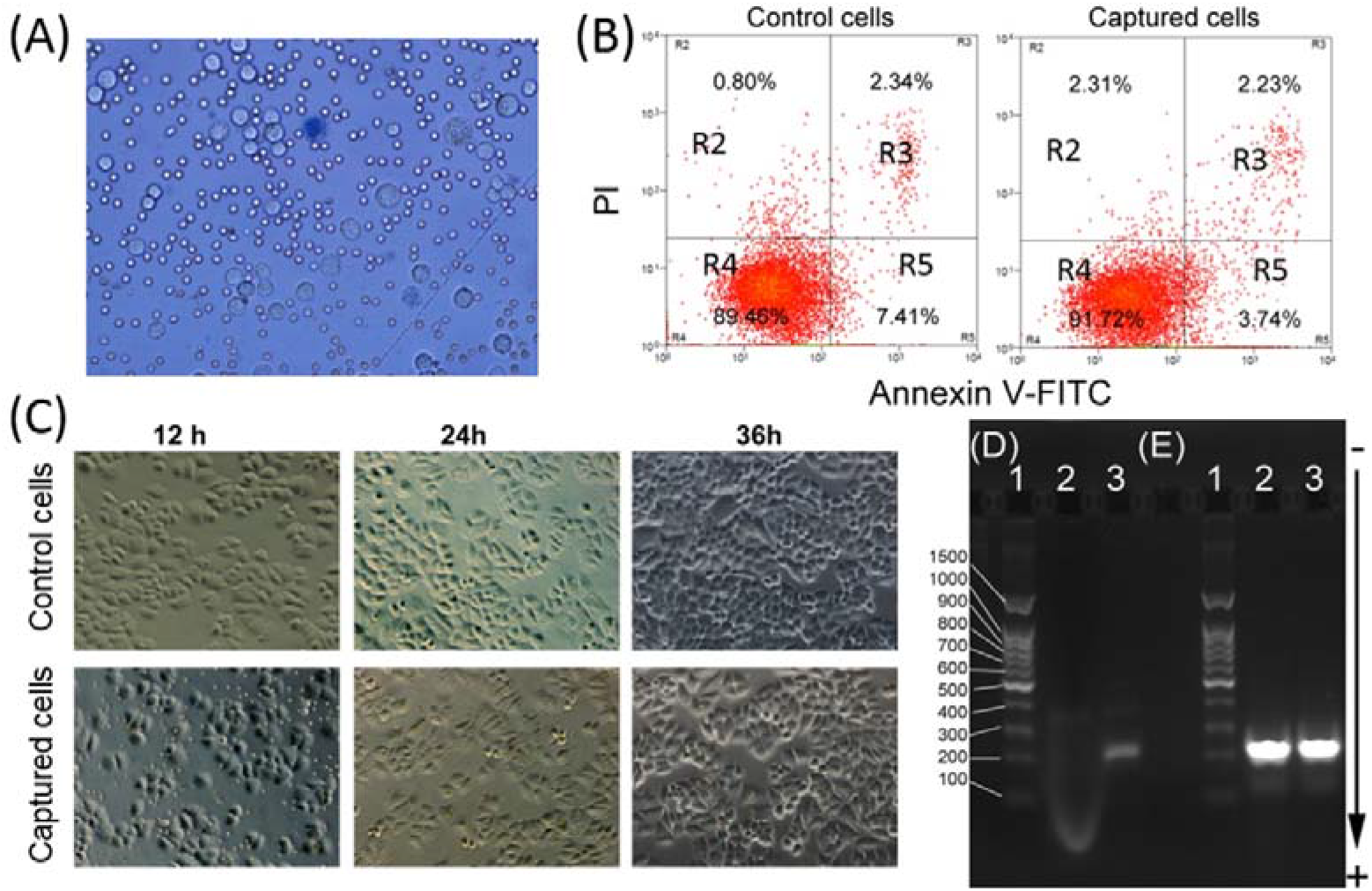
Viability and PCR analyses of the captured tumor cells. (A) Microscopic image of captured cells dyed by trypan blue. (B) Apoptosis analysis of the control of tumor cells and captured tumor cells by flow cytometry. At least 10,000 cells were measured per sample. The proportion (%) of cell number is shown in each quadrant. The proportion of viable cells was shown in the R4 quadrant (FITC-/PI-), early apoptotic cells shown in the R5 quadrant (FITC+/PI-), late apoptotic/necrotic cells shown in the R3 quadrant (FITC+/PI+). (C) Microscopic images of the control and captured tumor cells were plated and cultured for 12, 24, and 48 h. Agarose gel electrophoresis of products from RT-PCR amplification of EGFR mutation (D) and GAPDH (E). (Lane 1: DNA ladder, Lane 2: A549 cells captured with iFMNs, Lane 3: HCC827 cells captured with iFMNs)

Moreover, we also investigated the possibility of captured tumor cells for nucleic acid molecular analysis. Reverse transcriptase-PCR, which is widely used for precision tumor patient identification and target therapy, was performed to amplify epidermal growth factor receptor (EGFR) mRNA (Figure 6D) with Glyceraldehyde 3-phosphate Dehydrogenase (GAPDH) as a housekeeping gene (Figure 6E). Figure 6D shows that the 214-bp DNA band of the EGFR coding region was found with the HCC827 cells captured by iFMNS (lane 3 in Figure 6D), while no band was found when iFMNS were used to treat tumor cells (lane 2 in Figure 6D). These suggested iFMNS immunological binding had a negligible influence on RNA extraction and PCR reaction, and RT-PCR can analyze the isolated cells without disassociating iFMNS. Overall, the tumor cells captured with iFMNS had nearly no influence on tumor cells’ viability and were suitable for subsequent molecular biological analysis, which was crucial for further clinical diagnosis and research.

### CTC capture and testing in patient blood samples

To verify our method can be used for clinical samples, the peripheral blood from a lung cancer patient was collected, and the detection of CTC was performed in 2 days. As shown in Figure 7, intense red and blue fluorescence was observed from iFMNS and DAPI with a tumor cell. Meanwhile, no green fluorescence signal was observed in channel CD45-FITC. For captured cells that are CD45 positive and EpCAM negative with DAPI stained nucleus were identified as white blood cells. The results demonstrated that the as-prepared iFMNS could be used to simultaneously isolate and identify CTCs in real clinical samples with high efficiency.

**Figure 7.**
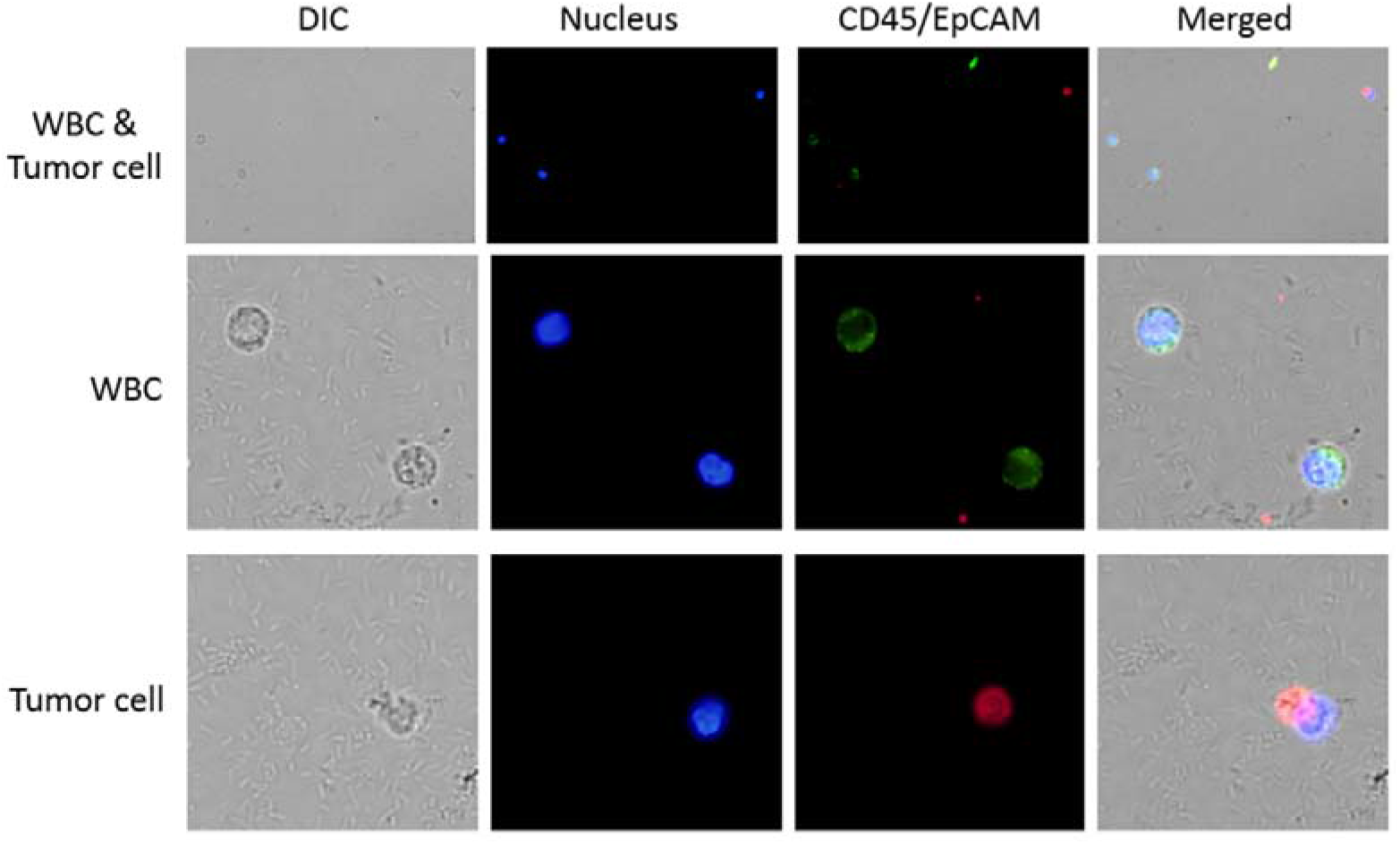
Images for enumeration of CTCs from a lung cancer patient by our method, CD45 antigen (green) was used to identify contaminating leukocytes, EpCAM antigen (red) was potentially used to identify CTCs.

## Conclusions

In summary, an ultra-bright fluorescent magnetic nanobeads system was successfully constructed for simultaneously efficient capture and sensitive detection of CTCs in whole blood by combining QDs, magnetic nanoparticles, and anti-EpCAM antibody. The developed FMN has several advantages that include simple and cost-effective preparation using emulsion/evaporation technique, ultra-bright fluorescence for sensitive target detection, easily separated by magnetic field for rapid target isolation, and biocompatible for antibody conjugation. With these exciting properties, iFMNS allows convenient and simultaneous tumor cell fluorescent immuno-labeling and magnetic isolation via the specific targeting to EpCAM. Compared with conventional immuno-magnetic isolation approaches, the fluorescent magnetic probes provide an additional signal for cell identification. The developed ultra-bright iFMNS are potential for rapid and simple CTCs isolation and can be adopted for multiplexed labeling to improve CTCs testing and analysis using QDs with different colors and fluorescence signals.

## Acknowledgments

We acknowledge research support from the International Science & Technology Cooperation Program of China (2014DFA33010) and Shanghai Strategic Emerging Industry Major Project (AI/RD2017-02).

